# Safety and Potency of COVIran Barekat Inactivated Vaccine Candidate for SARS-CoV-2: A Preclinical Study

**DOI:** 10.1101/2021.06.10.447951

**Authors:** Asghar Abdoli, Reza Aalizadeh, Hossein Aminianfar, Zahra Kianmehr, Ebrahim Azimi, Nabbi Emamipour, Hamidreza Jamshidi, Mohammadreza Hosseinpour, Mohammad Taqavian, Hasan Jalili

## Abstract

There is an urgent demand to manufacture an effective and safe vaccine to prevent SARS-CoV2 infection, which resulted in a global pandemic. In this study, we developed an inactivated whole-virus SARS-CoV-2 candidate vaccine named COVIran Barekat. Immunization at two different doses (3 µg or 5 µg per dose) elicited a high level of SARS-CoV-2 specific neutralizing antibodies in mice, rabbits, and non-human primates. The results show the safety profile in studied animals (include guinea pig, rabbit, mice, and monkeys). Rhesus macaques were immunized with the two-dose of 5 µg and 3 µg of the COVIran Barekat vaccine and showed highly efficient protection against 104 TCID50 of SARS-CoV-2 intratracheal challenge compared with the control group. These results highlight the COVIran Barekat vaccine as a potential candidate to induce a strong and potent immune response which may be a promising and feasible vaccine to protect against SARS-CoV2 infection.

## Introduction

The impact of vaccines on human health and control of infectious diseases is one of the brightest triumphs in the history of science.1 Candidate vaccine platforms for coronavirus disease in 2019 (COVID-19) are categorized into five major types: recombinant viral vectors, inactivated viruses, nucleic acid-based vaccines, protein subunit, and live attenuated virus.2 Inactivated vaccines are safe and effective since they cannot replicate at all in an immunized individual or there is no risk of reversion to a wild-type form which is capable of causing diseases.3

Currently, there are six licensed viral vaccines that are inactivated through either β-propiolactone (BPL) or formaldehyde. Beta-propiolactone is commonly used as an inactivating reagent for Rabies and Influenza virus vaccines whereas formaldehyde is used to inactivate Poliovirus (PV), Hepatitis A Virus (HAV), Japanese Encephalitis Virus (JEV), and Tick-Borne Encephalitis Virus (TBEV) in vaccine development.4 Initially, back in 1936, chemical inactivation was successfully applied to manufacture the Influenza vaccine.5 Experience with that vaccine led to a formalin-inactivated polio vaccine developed by Jonas Salk that came into use in 1955.6 Provost and coworkers prepared a hepatitis A vaccine based on chemical inactivation with formalin in 1986.7 The high efficacy of the hepatitis A vaccine testifies to the ability of careful inactivation to keep immunogenicity.

Vaccine development procedures depend on the selection of antigens, vaccine platforms, route of administration, and regimen number. Since the whole virus has all viral structural proteins, immune cells can recognize all viral immunogenic proteins present in its structure. An inactive SARS-CoV2 vaccine similar to the native virus has four structural proteins, known as the S (spike), E (envelope), M (membrane), and N (nucleocapsid) proteins.3 The S protein is the main target for neutralizing antibodies in all coronaviruses, which is composed of S1 and S2 domains. SARS-CoV-2 spike protein covers several distinct antigenic sites, comprising several receptor-binding domain (RBD) epitopes along with non-RBD epitopes.8 Levels of antibodies rise against nucleoprotein (N) which is the most abundant viral protein.9,10 Although N antibodies are unlikely to neutralize the virus, they have been indicated to confer protection against mouse hepatitis virus and coronavirus in mice.11

The duration of the antibody response for SARS-CoV2 has not been well understood. However, previous longitudinal studies of patients with SARS-CoV infection have shown that neutralizing SARS-CoV antibodies last more than 3 years after the onset of symptoms (12). Inactivated vaccines are formulated with potent adjuvant, and need boosters to induce satisfactory and long-term immunity. The most commonly used adjuvants in human vaccines are aluminum salts and their activity was defined in 1926.13 Due to COVID-19 cytokine storm, alum, an adjuvant that promotes TH2-type immunity, actually reduces immunopathology and seems to be a suitable option for vaccine formulation.14

Inactivated SARS-CoV2 vaccines may have better efficacy against variants that have mutations in the spike protein. In the current study, the efficiency and safety of an inactivated whole-virus SARS-CoV-2 candidate vaccine (COVIran Barekat vaccine) were evaluated in a preclinical trial.

## Materials and Method

### Virus Isolation and Characterization

All experiments with live SARS-CoV-2 viruses were performed in a biosafety level-3 facility according to WHO guidelines.15 The virus was isolated from the nasopharyngeal sample from a COVID-19 patient. Before being infected with monolayer Vero cells (ATCC# CCL81), samples were diluted with a viral transfer medium containing antibiotics and passed through a 0.22 µm filter (Sartorius, Germany). Infected Vero cells were maintained in Dulbecco’s modified Eagle’s medium (Bio Idea, Iran) supplemented with 10% heat-inactivated fetal bovine serum (FBS) and incubated for 72 h at 37°C in a 5% CO_2_ incubator. Virus growth was verified through cytopathic effects (CPE), gene detection, and electron microscopy.

The SARS-CoV-2 virus titer was determined by 50% tissue culture infective dose (TCID50). Vero cells (1–2 × 10^4^ cells/mL) were seeded in 96-well plates and incubated for 16–24 h at 37°C. Serial 10-fold dilutions of virus-containing samples were added to the 96-well culture plate and cultured for 3–7 days in a 5% CO_2_ incubator at 37□, and cells were checked under a microscope for the presence of CPE. The virus titer was calculated by the Spearman Karber method.16

After 4 passages, viral RNA was extracted from the clarified cell culture supernatant of infected Vero cells by RNA extraction kit (Rojetechnologies, Yazd, Iran), and converted to cDNA. Whole-genome sequencing was carried out using the Sanger method. To evaluate the genetic stability of the virus, five more passages were prepared to obtain the P10 stock and the whole genome was sequenced again.

### Vaccine Preparation

#### Virus Inactivation and its Validation

SARS-CoV-2 virus culture supernatant was harvested 72 h post-infection and clarified. After that, the virus particles were inactivated using β-propiolactone at 2–8°C for 20–24 h. To confirm the effectiveness of the virus inactivation procedure, the inactivated SARS-CoV-2 virus was inoculated onto Vero monolayers and incubated at 37°C in a 5% CO_2_ incubator. Daily CPE monitoring was performed consecutively for three blind passages under a microscope. No CPE was observed in three passages, while positive controls showed 100% CPE.

#### Purification

The inactivated virus solution was concentrated via ultrafiltration and buffer was exchanged using diafiltration. Purification of the inactivated virus was done by ultracentrifugation and chromatography methods. The protein composition of the purified inactivated virus was confirmed through SDS-PAGE analysis followed by staining with silver nitrate and western blot. The total protein concentration was determined by the Bradford assay.

The structural integrity of the isolated virus particles was monitored using the transmission electron microscope (TEM). Viral particles were absorbed onto carbon-coated grids and stained with 1% uranyl acetate. Grids were analyzed using a transmission electron microscope (TEM, EM208S, Philips, 100 KV) at final magnifications of 90000×.

#### Formulation

The purified inactivated viral particles (antigen) were sterilized using the filtration, and a mixture 2% adjuvant® Alhydrogel-antigen in two doses, a high dose of 5µg and a low dose of 3µg, of inactivated viral antigen was prepared. To improve efficient absorption of antigen, 2% adjuvant® Alhydrogel (Brenntag Biosector, Denmark) was used for well mixing.

### Animal Studies

All experiments with animals (BALB/c mice, guinea pigs, rabbits, and macaques) were carried out in accordance with the approval of the animal ethics committee guidelines of the Ministry of Health and Medical Education (Tehran, Iran). The schematic diagram of animal studies on the COVIran Barekat vaccine is shown in Figure 1.

**Figure 1.**
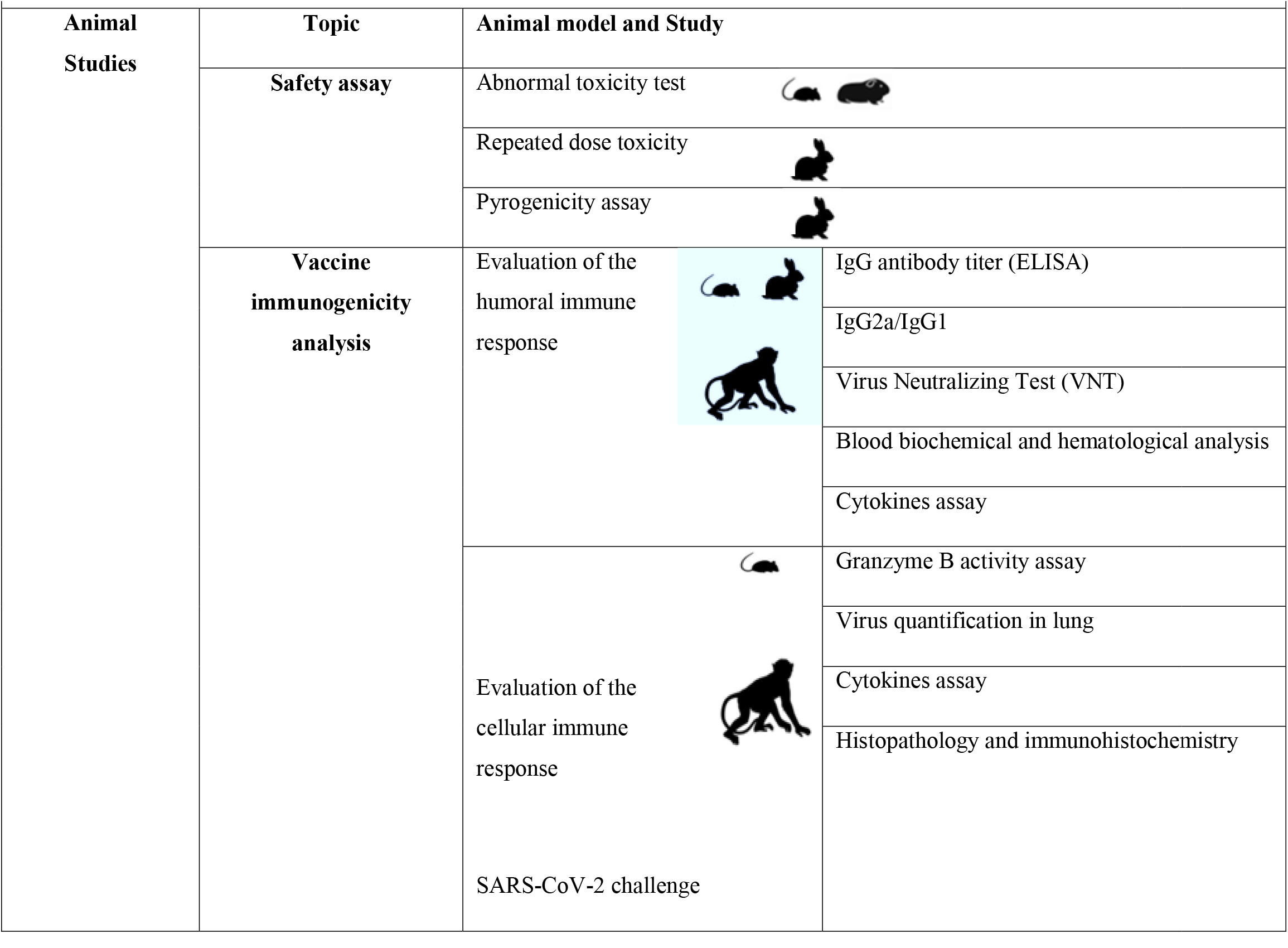
Animal studies for evaluating the safety and efficacy of the COVIran Barekat vaccine.

#### Safety Assay

In order to evaluate abnormal toxicity, 7 guinea pigs and 5 mice were injected intraperitoneally with 5 µg antigen (equivalent to the human dose) in combination with Alum adjuvant, and 3 guinea pigs and 2 mice as control group were injected with water for injection (WFI), as well. The clinical signs, mortality, and changes in body weight were monitored in all animals for up to 14 days. The site of injection was also evaluated for erythema and edema at 24, 48, and 72 hours to check for local tolerance. To study toxicity in repeated doses, the rabbits were injected with 5µg of the vaccine up to 3 times. All animals were examined for mortality during the experimental period and were euthanized, necropsied, and the organs were dissected for macroscopic and microscopic evaluation.

To conduct the pyrogenicity assay, nine rabbits with approximately the same weight were selected for 3 consecutive batches of candidate vaccines. Rabbits were deprived of food and only had access to water 24 hours before the injection. After one hour, the initial temperature of the animal was recorded by a special rectal thermometer placed in the animal’s rectum. Five μg of the candidate vaccine was injected into the marginal vein of the rabbit’s ear. The post-injection body temperature of rabbits was measured in the first, second, and third hours using a special thermometer.

#### Vaccine Immunogenicity Analysis

Two doses of the inactivated purified virus (3 and 5 µg) as antigen alone or in combination with alum adjuvant were administrated to different animals to evaluate vaccine immunogenicity. The control group was injected with physiological saline (PBS).

##### Mice

1) Ten female BALB/c mice (6–8 weeks) were injected intramuscularly with 100 μL of different doses of the vaccine according to Table 1 on days 0 and 14. Two and five weeks after the final immunization, blood samples were collected by puncture of the retro-orbital vein and serum samples were taken to evaluate the antigen-specific antibody binding titer and antibody isotyping profile by Enzyme-Linked Immunosorbent Assay (ELISA), and virus-neutralizing assay (VNT) was applied to analyze the vaccine’s immunogenicity.

**Table 1.**
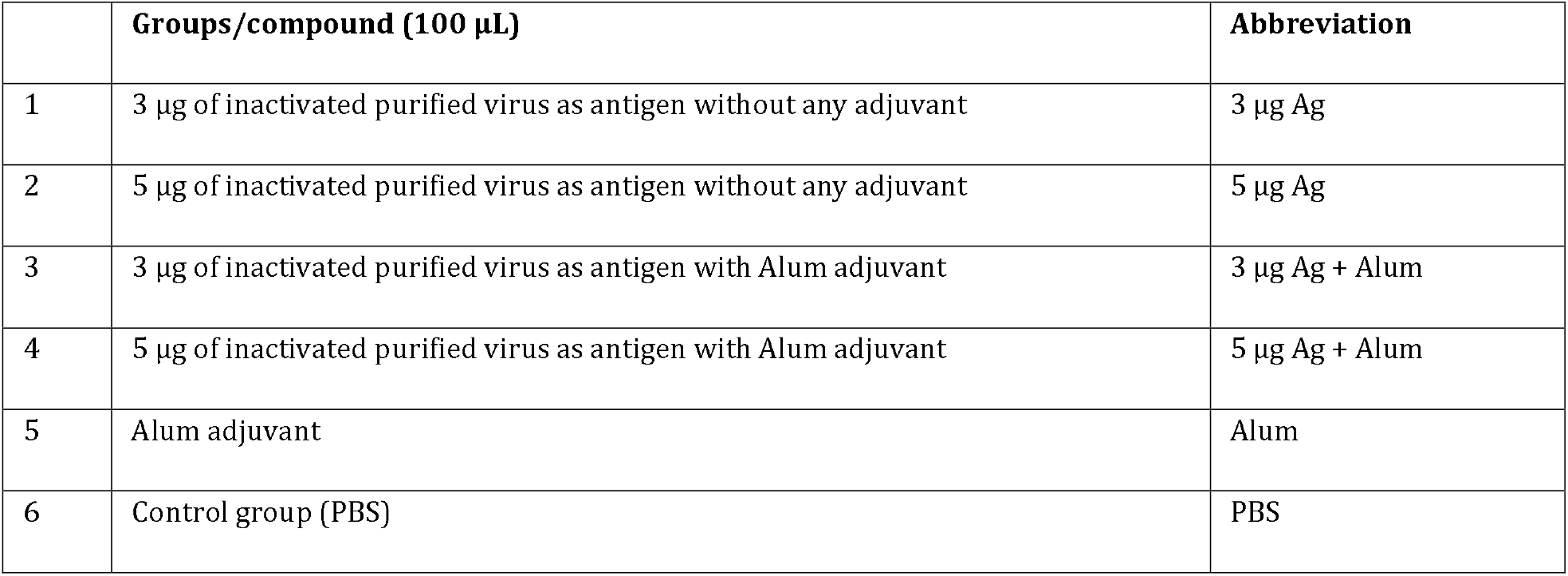
Different compounds used for injection in the first mice groups

|  | Groups/compound (100 µL) | Abbreviation |
| --- | --- | --- |
| 1 | 3 µg of inactivated purified virus as antigen without any adjuvant | 3 µg Ag |
| 2 | 5 µg of inactivated purified virus as antigen without any adjuvant | 5 µg Ag |
| 3 | 3 µg of inactivated purified virus as antigen with Alum adjuvant | 3 µg Ag + Alum |
| 4 | 5 µg of inactivated purified virus as antigen with Alum adjuvant | 5 µg Ag + Alum |
| 5 | Alum adjuvant | Alum |
| 6 | Control group (PBS) | PBS |

2) Ten BALB/c mice were injected with the candidate vaccine using the same protocol presented in Table 1. Two weeks after the final immunization, serum samples were collected to evaluate the vaccine’s immunogenicity by virus-neutralizing assay. Then, the animals were sacrificed, spleens were isolated and splenocyte culture supernatants were used for evaluating cellular immunity responses by Granzyme B activity assay. Rabbit: Fifty Dutch-Polish rabbits (3–6 months) were divided into two groups: the test and control groups. The test group was injected intramuscularly with 5 µg antigen in combination with alum adjuvant and the control group was injected with physiological saline (PBS). Immunizations were carried out 3 times (on days 0, 14, 28) using 5 μg of antigens in combination with alum. After the third injections, the rabbits were bled and individual sera were assayed at 7 and 28 days after the last injection.

##### Monkey

Rhesus macaques (3– 4 years old) were divided into three groups: in the placebo group, the animals were injected intramuscularly with physiological saline (PBS), and the two test groups were injected intramuscularly with 3 and 5 µg antigen in combination with alum adjuvant. All macaques were vaccinated on days 0 and 14. A challenge study was carried out 10 days after the second immunization under anesthesia through direct intratracheal inoculation of 10^4^ TCID50 of SARS-CoV-2. Safety parameters including immunization evaluation before and after the challenge, daily clinical and general observation, body weight, and body temperature were collected and preserved as a database. Peripheral blood was collected on days 0, 7, 14, and 21 post-immunizations and one week after the challenge. Analysis of S-specific IgG antibody titer, biochemical blood test and hematological indices was performed in collected blood samples.

Euthanasia was performed by intravenous administration of sodium thiopental (100 mg/kg), followed by complete systematic necropsy and sampling. In addition to macroscopic evaluation of organs and the site of injection, tissue sampling was performed for histopathology. Lung tissue samples were quickly frozen for cytokine evaluation and some other samples by qRT-PCR.

All animals were euthanized on day 14 post-challenge, and different organs such as the lung, heart, spleen, liver, kidney, and brain were collected. Lung tissues were used for measurement of the viral load (RT-PCR assay) and several cytokine levels were determined via western blotting, histopathology pattern determination and characterization of lung immune cell population by immunohistochemistry.17,18

## 1. Evaluation of the Humoral Immune Response

### a) SARS-CoV-2 Specific IgG Antibody Titer Assay by ELISA

The SARS-CoV-2 specific IgG antibody titer of collected serum samples from all immunized animals was determined through enzyme-linked immunosorbent assay (ELISA). A 96-well plate was coated with the purified whole inactivated virus solution (5 μg/mL) in phosphate-buffered saline (PBS, pH 7.2) by overnight incubation at 4°C. Then, the plates were washed three times with PBS containing 0.05% Tween 20 (PBST) and blocked with 5% skim milk in PBS at 37°C for 1.5 h. After the washing step, serial dilutions of sera in PBS were added to the plates and incubated at 37°C for 1.5 h. Horseradish peroxidase-conjugated goat anti-mouse IgG antibody (RaziBiotec AP8036) at 1/5,000 dilution was added to the wells (goat anti-rabbit (Abcam) at 1/20,000 dilution used in ELISA of rabbit serum) and after washing with PBST, the plates were incubated for 1.5 h at 37°C. Colorimetric detection was performed using 100μL tetramethylbenzidine (TMB, Sigma) as substrate after washing and incubation for 30 min at RT. The reaction was stopped with 50 μL H2SO4 (2 N), and absorbance was measured at 450 nm using an ELISA reader.

Furthermore, N and S-specific IgG antibody titers in serum samples of the monkey group were determined by Covid-19 IgG ELISA Kit (Padtangostar, Iran). Briefly, dilutions of 1/100 sera were added to the plates coated with the purified N and S proteins and incubated at 37°C for 30 min. The plate was washed three times with washing buffer and then, goat anti-monkey IgG antibody conjugated with HRP (Abcam, 190549,) at 1/100 dilution was added to the wells and the plate was incubated for 30 min at 37°C. Following the washing step, colorimetric detection was performed using TMB as the substrate for 15 min at RT. The reaction was stopped with 50 μL H2SO4 (2 N), and absorbance was measured at 450 nm using an ELISA reader.

#### b) Immunoglobulin (IgG) Subclass (IgG2a/IgG1 Assessment)

In this study, 96-well ELISA Maxisorp (96-well) plates coated overnight at 37°C with 50 μL of SARS-CoV-2 (whole inactivated antigen) protein were washed with PBS containing 0.05% Tween 20, retained for 1 h at 37°C with 1% bovine serum albumin (BSA) solution made in 1X PBS/Tween 20, and then washed with (PBS-0.05% Tween 20). Sera were added to each well and incubated at 37°C for 2 h. The plates were washed with PBS-Tween 20 (0.1%). After that, horseradish peroxidase-conjugated goat anti-mouse IgG1 and IgG2a were added to the plate and incubated for 1 h at 37°C. The wells were washed and 100 µL of TMB substrate was added and incubated for 30 min. After stopping the reaction, absorbance was read at 450 nm by Universal Microplate Reader (Biohit 8000, USA).

#### c) Virus Neutralizing Test

The virus neutralization test (VNT) was performed to analyze vaccine protectivity. Briefly, 50 µL of two-fold serial diluted sera were mixed with 50 µL of 100 TCID50/mL of SARS-CoV2 in DMEM and incubated for 60 min at 37°C. The virus/serum mixtures were then inoculated onto monolayers of 1–2×10^4^ Vero cells in 96-well plates to attach free viruses for 60 min at 37°C. The supernatant was removed at 1 h post-infection, the infected cells were washed twice with DMEM and kept in DMEM for 48 h at 37°C in a 5% CO_2_ incubator. The cytopathic effect (CPE) of each well was recorded under microscopes, and the neutralizing titer was calculated by the dilution number of 50% protective condition.19,20

#### d) Blood Biochemical and Hematological Analysis in Rhesus Macaques

Blood sampling and evaluations were performed weekly. Biochemical markers including glucose, urea, creatinine (Cr), total protein, albumin, C-reactive proteins (CRP), alanine aminotransferase (ALT), aspartate aminotransferase (AST), and lactate dehydrogenase (LDH) were analyzed by an autoanalyzer using animal specific kits. Hematology including measurement of White blood cell (WBC), Red blood cell (RBC), Neutrophil, Lymphocyte, Monocyte and Eosinophil count, hemoglobin (Hb), hematocrit (HCT), mean corpuscular volume (MCV), mean corpuscular hemoglobin (MCH), mean corpuscular hemoglobin concentration (MCHC), red cell distribution width (RDW) and platelet count (PLT) were performed by a cell counter (Nihon Kohden Celltac alpha MEK-6450 hematology analyzers).

#### e) Lung Tissue Cytokine Assessment of Rhesus Macaques

The lung tissue samples were used for protein extraction with RIPA buffer containing 50 mM Tris HCl pH 7.4, 150 mM NaCl, 5 mM EDTA, 1% Nonidet P-40, 1% sodium deoxycholate, 0.1% SDS, 1% aprotinin, 50 mM NaF and 0.1 mM Na3VO4 (CitoMatin Gene) under class II biosafety cabinet.

The total protein concentration was determined by the Bradford assay. The assessment of interferon (IFN)-γ, and interleukin (IL)-6, IL-4, IL-10 in lung extracted proteins was performed through SDS-PAGE (Padideh Nojen UV-80V system with a standard protocol) followed by western blot analysis using anti-IL-4 (ab34277), anti-IL-6 (ab233706), anti-IL-10 (ab133575), anti-IFN-γ (ab231036) and anti-β actin as normalized control (ab170325) according to the standard protocol. Cytokine levels were normalized to the total protein concentration in each lung homogenate. Western blot images were analyzed using NIH Image J computer software and the relative intensities of each cytokine bands to β-actin protein were calculated.

## 2. Evaluation of the Cellular Immune Response

### a) Granzyme B Activity Assay in Mice

The potential of cellular immunity induction was evaluated by Granzyme B (Gzm B) activity in different vaccine preparations. Spleens were isolated from the immunized mice 2 weeks post-immunization, and single-cell suspensions were prepared by gentle homogenization. Red blood cells (RBCs) were lysed by incubation in RBC lysis buffer (20 mM Tris, 160 mM NH4Cl, pH 7.4) for 5 min at RT, and the resulting splenocytes were resuspended at a density of 2 × 10^6^ cells/mL in RPMI 1640 supplemented with 10% FBS, 2 mM L-glutamine, 100 U/mL penicillin and 100 μg/mL streptomycin after washing twice with RPMI. Cells were seeded in a flat-bottom 24-well plate (Nunc, Denmark) and stimulated with purified inactive whole SARS-CoV-2 particles and then incubated at 37°C with 5% CO_2_. Culture supernatants were collected for measurement of Gzm B activity after 72 h. Gzm B activity was determined using commercial mouse Granzyme B ELISA Ready-SET-Go kit (88-8022, eBioscience).

### b) SARS-CoV-2 Quantification in Lungs of Challenged Rhesus Macaques

To remove cell debris, lung homogenates were clarified via centrifugation at 4000 **g** for 10 min, and the supernatants were then stored at −80°C until the assay was carried out. Vero cells with 80 confluency in 96-well plates were infected with serial log10 dilutions of clarified lung samples in DMEM. After 72 h of incubation at 37°C in a 5% CO_2_ atmosphere, the culture supernatants were tested for the presence of SARS-CoV2. The viral titer in each lung specimen was calculated by the Karber method and expressed as the 50% tissue culture infective dose (TCID50).

### c) Histopathology and Immunohistochemistry in Rhesus Macaques

Tissues were fixed in formalin and embedded in paraffin blocks. 5 µm sections were stained with hematoxylin and eosin for histopathological interpretation. 3 µm sections of lung paraffin blocks were used for immunohistochemical staining. Anti-CD4 (CMGCD4-S50), Anti-CD8 (BRB036-ZYTOMED), Anti-CD20 (BMS003-ZYTOMED), and Anti CD68 (MAD-002097QD) were the markers which were applied for detection of immune cells in different parts of lung sections. Throughout each slide, 10 randomly chosen fields were evaluated in a high power field (HPF) for counting CD4, CD8, CD20 lymphocytes, and CD68 positive macrophages by light microscopy (Nikon Eclipse E600, Japan).

### 2.4. Statistical Analysis

Analysis of different data was performed using one-way ANOVA or two-way ANOVA followed by Tukey’s post hoc test. Values were expressed as mean ± SD and *P* < 0.01 was considered as statistically significant. All statistical analyses were performed using the GraphPad Prism 7 software (La Jolla, CA, USA).

## Results

### Virus Isolation and Characterization

The SARS-CoV-2 used in developing the COVIran Barekat vaccine was retrieved from a nasopharyngeal sample from a COVID-19 patient. The sample propagation and virus isolation were performed in the Vero CCL-81.

For efficient growth of viral stock in Vero cells, the isolated virus was first plaque purified and passaged once in Vero cells to generate the P1 stock. After this, three other passages were performed to generate the P2 to P4 stocks. At multiplicities of infection (MOI) of 0.001-0.01, multiplication kinetics analysis of the P4 stock in Vero cells showed that the stock replicated efficiently and yielded a peak titer of 6 log10 TCID50/mL by 3 days after infection (dpi). After 4 passages, the master and working vaccine seed was prepared. Virus growth and isolation were verified through cytopathic effects and gene detection. In terms of the overall divergence of isolated SARS-CoV-2, this strain had 99.9% identity to the earliest detected strain, Wuhan Hu-1.21 Results showed genetic stability of the isolated virus in multiple passages of Vero cells which is suitable for vaccine development. The flowchart of COVIran Barekat vaccin preparation is illustrated in Figure 2.

**Figure 2.**
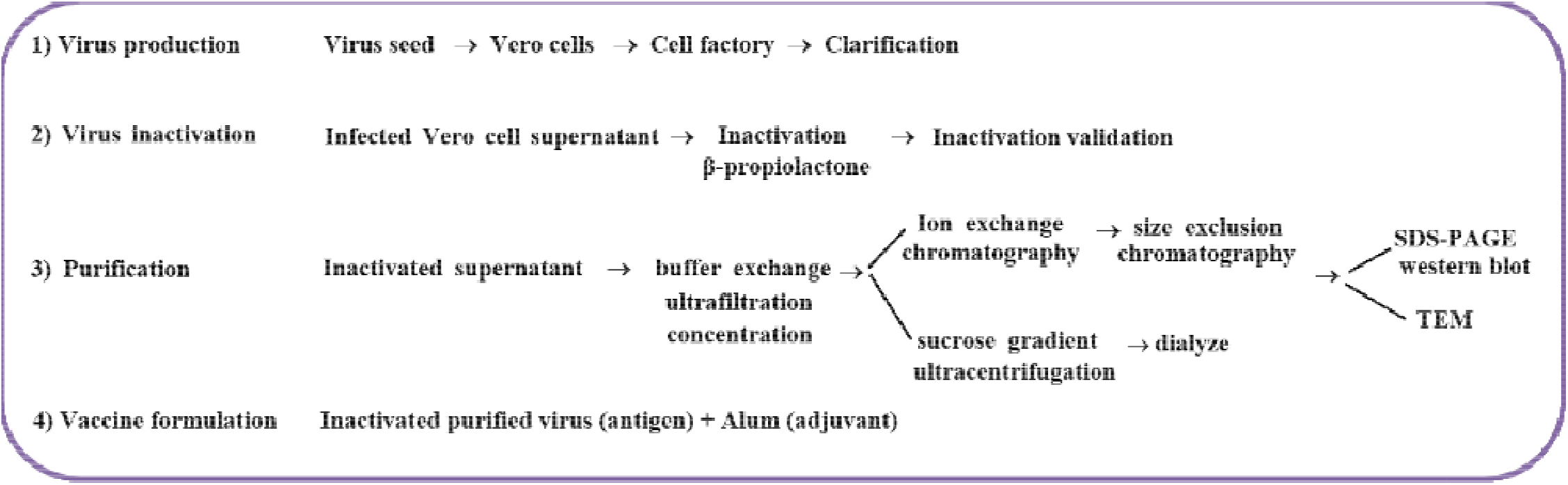
Flowchart of COVIran Barekat vaccine preparation.

Growth kinetics analysis showed that the virus replicated to 7.0 log10 TCID50 at 72 h post-infection. The infected Vero cell culture supernatant was harvested 72 h post-infection and viral particles were inactivated via β-propiolactone. To verify the inactivation process, the third blind passage was carried out for three generations from the inactivated virus and did not show any CPE (Figures 3A, 3B, and 3C). The inactivated virus was concentrated through ultrafiltration and purified using ion-exchange and size exclusion chromatography or the discontinuous sucrose gradient (20/60% w/v) ultracentrifugation.

**Figure 3.**
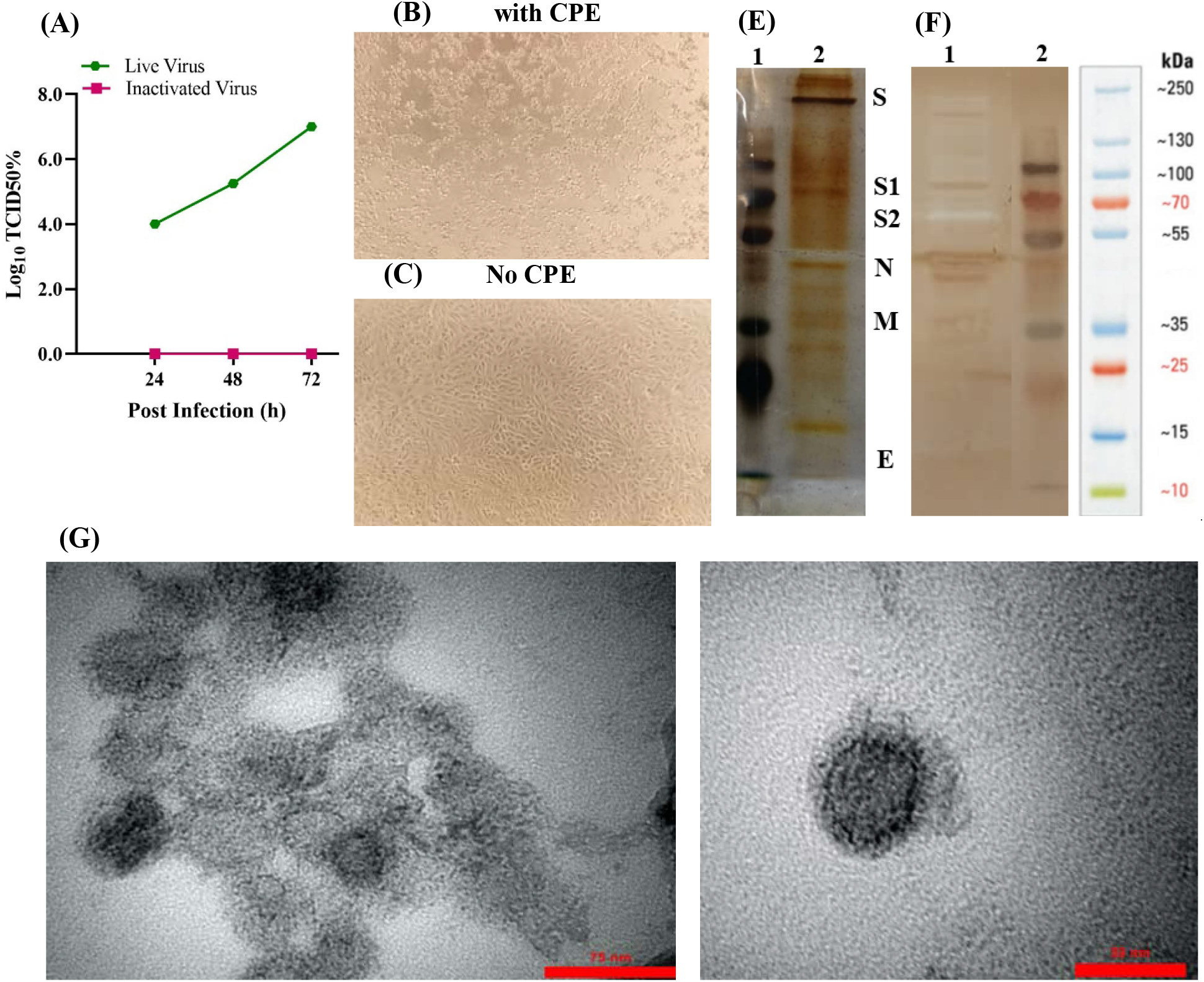
Analysis of the purified inactivated viral particles. (**A**) Cytopathic effect (CPE) of SARS-CoV-2 virus before and after inactivation. Virus titer (10 -10) measured by CCID50 at three time-points: 24, 48, and 72 h. (**B**) Image of Vero cell monolayer with CPE before inactivation and (**C**) no CPE after inactivation. (**E**) Samples of the purified inactivated viral particles were separated on 10% SDS-PAGE and stained with silver nitrate: 1, molecular size markers; 2 purified viral particles. (**F**) Western blot: proteins on SDS-PAGE gel were transferred onto PVDF membrane and SARS-CoV-2 proteins were detected using the anti-rabbit polyclonal antibody: 1, purified viral particles and 2, molecular size markers. SDS-PAGE and Western blot patterns show the major structural proteins: spike protein (**S**), nucleocapsid protein (**N**), membrane protein (**M**), and envelope protein (**E**). (**G**) Electron micrographs of negatively stained purified viral particles under TEM. Purified viral particles were negatively stained with 1% uranyl acetate and observed under TEM (TEM, EM208S, Philips, 100 KV) at 90,000 × magnification.

Four major structural proteins with their corresponding equivalent molecular weights were identified by protein composition analysis of the purified inactivated virus particles using SDS-PAGE and western blot: the spike (S) protein, nucleocapsid (N) protein, membrane (M) protein, and the envelope (E) protein (Figures 3E and 3F). Transmission electron microscopy (TEM) analysis showed that the purified inactivated viral particles were intact, oval-shaped, and accompanied by a crown-like structure representing the well-defined spike on the virus membrane (Figure 3G). These results showed that the final purified inactivated bulk of the candidate vaccine is highly pure and contains the majority of structural proteins.

### Safety Assay

The purified inactivated viral antigen was formulated with Alum adjuvant and safety studies were performed on 7 guinea pigs which were injected with the 5 µg human equivalent dose intraperitoneally. The results of this test were monitored for 14 days as shown in Table 2. No pathological symptoms (gross lesions) or weight loss were seen in any of the vaccinated pigs and mice. Microscopic evaluation of tissue samples (liver, kidney, heart, lung) found no toxic changes due to vaccine administrations. To evaluate toxicity in the repeated doses, ten rabbits were injected with 5 µg of the vaccine up to 3 times. Laboratory results showed no mortality or pathogenicity in any of the groups and also the average weight increased from 1900 g to 4500 g 90 days post-infection.

**Table 2.** Safety evaluation of vaccinated guinea pigs and mice.

| Groups | Viability | Illness | Gross lesions | Weight loss |
| --- | --- | --- | --- | --- |
| **Purified inactivated virus (5 µg) + Alum**<br>Guinea pigs ($n = 7$) and Mice ($n = 5$) | yes | no | no | no |
| **Control (D.W)**<br>Guinea pigs ($n = 3$) and Mice ($n = 2$) | yes | no | no | no |

The results of the pyrogenicity assay in rabbits (Table 3) showed that the total increase in body temperature of the three rabbits at 3 different intervals was below the threshold (>1.2°C). Also,the difference between the body temperature of each animal at zero hours, and the following hours was up to 0.4°C. Therefore, it can be concluded that the candidate vaccine is not pyrogenic.

**Table 3.** Rabbit body temperature (°C) after fever test.

| **Table 3.** Rabbit body temperature (°C) after fever test. |  |  |  |
| --- | --- | --- | --- |
| **Initial temperature** | **after one hour** | **after two hours** | **after three hours** |
| 38.2 | 38.4 | 38.5 | 38.5 |
| 38 | 38.2 | 38.4 | 38.4 |
| 38.5 | 38.6 | 38.7 | 38.7 |

### Immunogenicity Assay

BALB/c mice, Dutch-Polish rabbits, and monkeys were immunized at two times (on days 0 and 14) with two doses of 3 or 5 µg/dose of the candidate vaccine, and the control group was injected with PBS. No inflammation or other adverse effects were observed. To analyze the vaccine’s immunogenicity, the humoral and cellular immune responses were evaluated by different methods including specific IgG antibody titers and IgG2a/IgG1 assessment (ELISA), virus-neutralizing assay and granzyme B activity assay in serum samples, biochemical and hematological analysis in peripheral blood samples, cytokines assay (western blotting), histopathology and immunohistochemistry in lung tissue.

## 1. Evaluation of the Humoral Immune Response

### a) Antibody Assay and VNT

Increasing serum anti-SARS-CoV-2 IgG antibody titers in vaccinated mice was monitored 2 and 5 weeks post-immunization using indirect ELISA. The results showed that SARS-CoV-2 specific IgG quickly boosted in the sera of vaccinated mice, and the titer of COV-2 specific IgG was significantly (*P* < 0.001) higher in the mice immunized with a combination of the purified inactivated viral particles (antigen) and alum adjuvant compared to the mice administered with antigen alone. Furthermore, in all groups administered with antigen alone or in combination with adjuvants, there was a significant increase of antibody titer (*P* < 0.0001) in groups that received the 5 μg dose of antigen in comparison with groups immunized with a dose of 3 μg antigen (Figure 4). In addition, IgG1 and IgG2a antibodies were raised in all mice that received the alum formulated vaccine (Figure 5).

**Figure 4.**
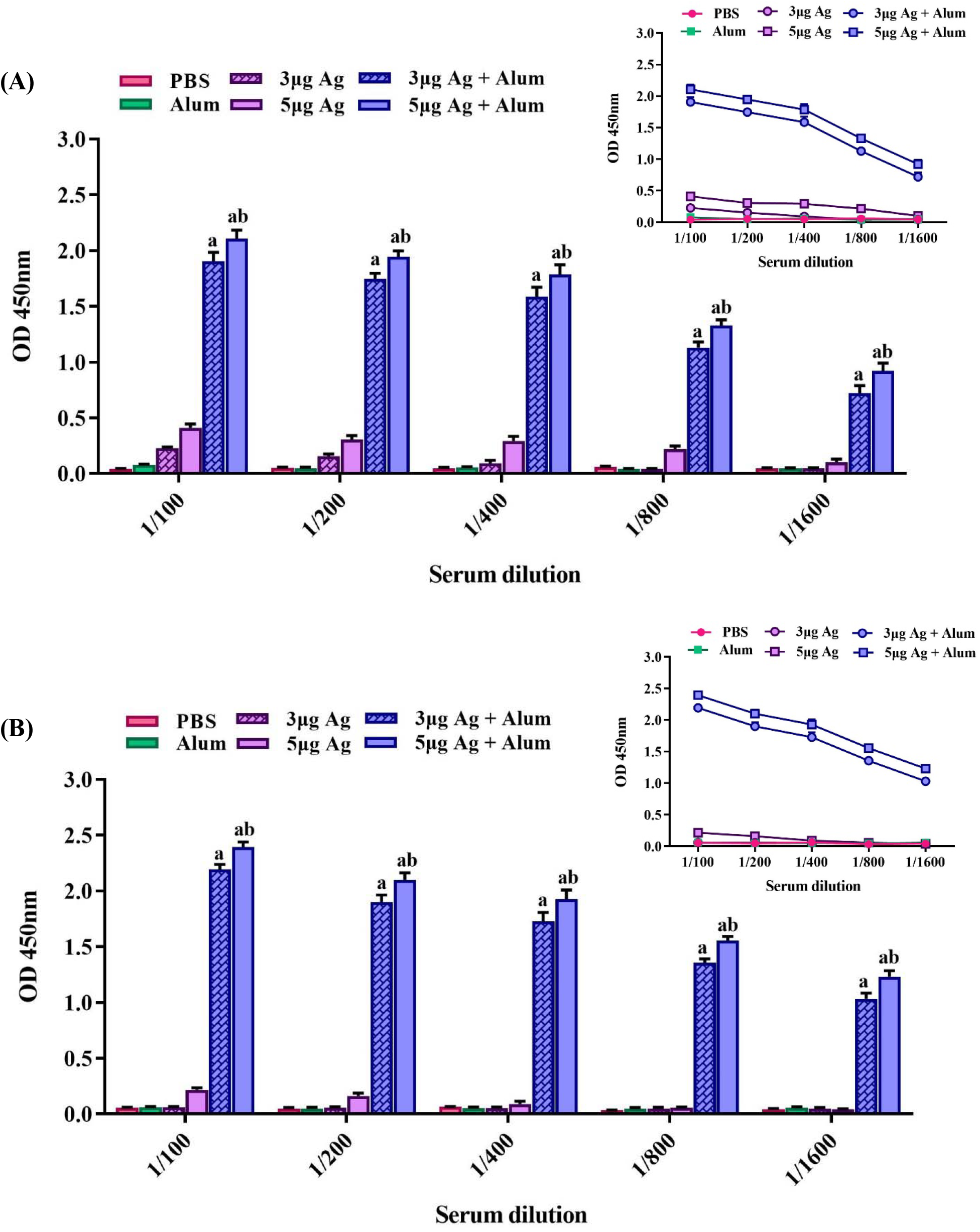
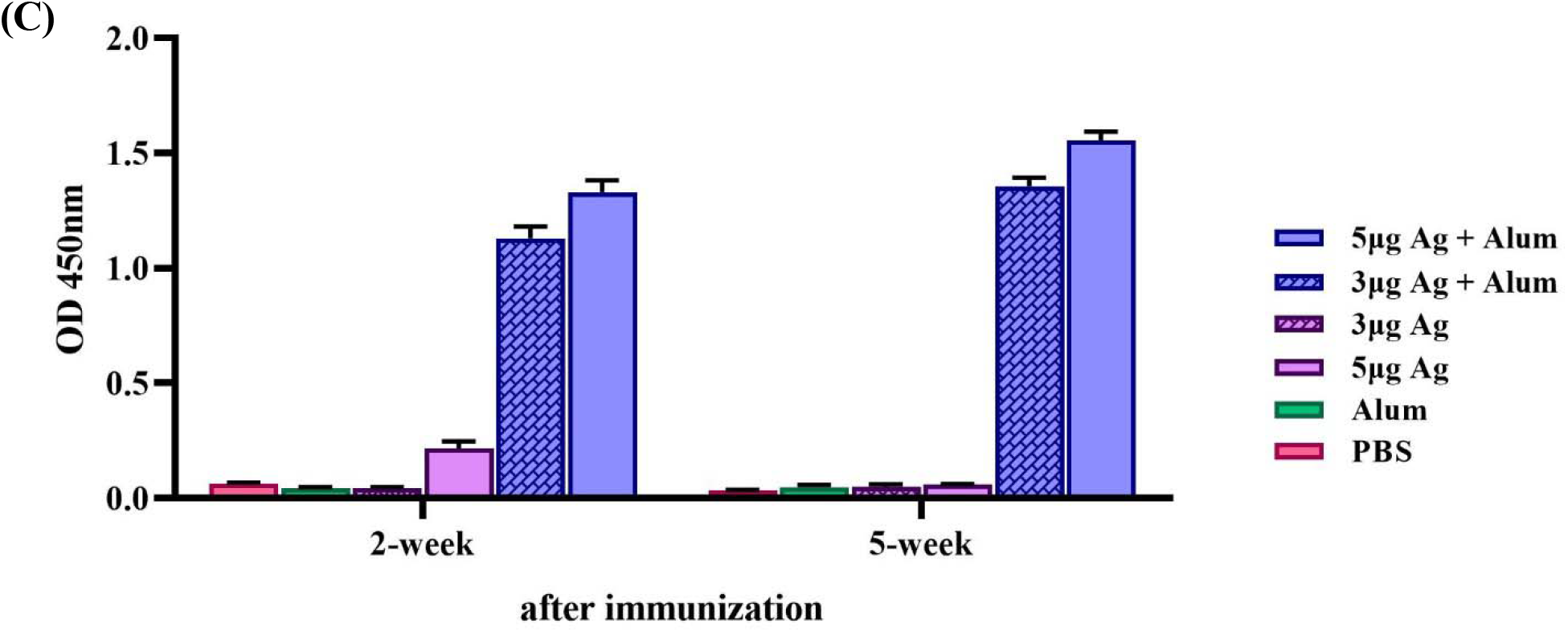
Induction of specific IgG antibodies by different vaccine preparation in mice groups. Immunizations were carried out two times during 2 weeks with 100 μL intramuscular injection of different vaccine preparations according to Table 1: a PBS solution containing 3 or 5 μg of purified inactivated viruses as antigens, administered alone or in combination with alum. The control groups received alum or PBS. At 2 and 5 weeks post-immunization, the mice were bled and individual sera were assayed by ELISA. Anti-COV IgG antibody titers in different dilutions of serum at 2 weeks post-immunization (**A**), 5 weeks post-immunization (**B**), and in 1/800 dilutions of serum (**C**). Data are mean ± SD. The levels of statistical significance for differences between test groups were determined using two-way ANOVA followed by Tukey’s pos thoc test. a: Statistical significance (P < 0.0001) in comparison with groups that received 3 or 5 μg antigen alone and b: Statistical significance (P < 0.0001) between groups that received 3 μg antigens and groups immunized with 5 μg antigens.

**Figure 5.**
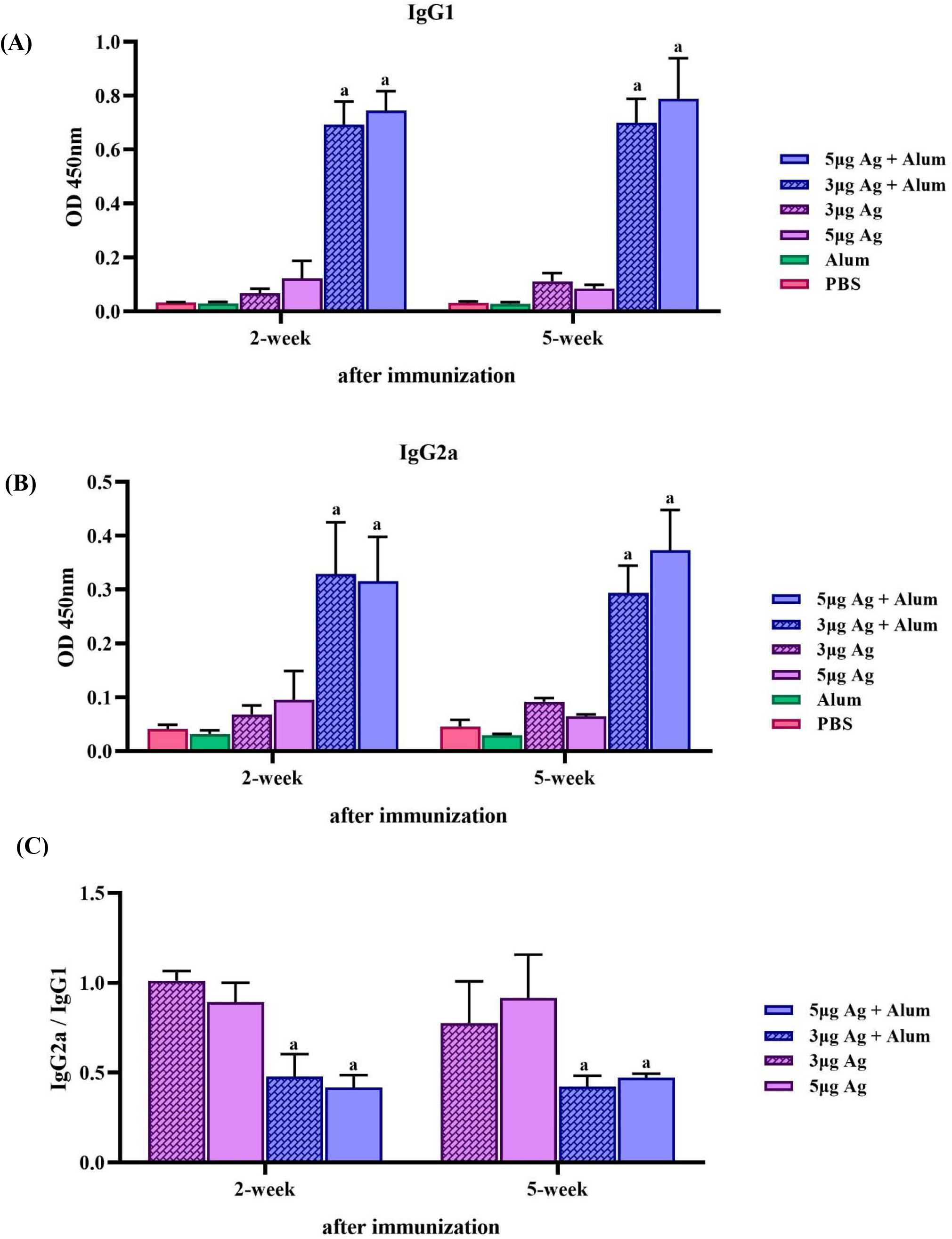
Measurement of specific IgG1, IgG2a antibodies and IgG2a/IgG1 ratio in mice groups that received different vaccine preparations. Immunizations were carried out two times during 2 weeks with 100 μL intramuscular injection of different vaccine preparations according to Table 1: a PBS solution containing 3 or 5 μg of purified inactivated viruses as antigens, administered alone or in combination with alum. The control groups received alum or PBS. The mice were bled at 2 and 5 weeks post-immunization, and individual sera were assayed by ELISA. (**A**) Anti-COV IgG1, (**B**) IgG2a antibody titers in 1/1000 dilutions of serum at 2 and weeks post-immunization and (**C**) IgG2a/IgG1 ratio. Data are mean ± SD. The levels of statistical significance for differences between test groups were determined using two-way ANOVA followed by Tukey’s post hoc test. a: Statistical significance (P 0.01) in comparison with groups that received 3 or 5 μg antigen alone.

It was found that the formulated vaccine (antigen + alum) increased the antibody titer after 42 days due to antigen storage, while antibody titers decreased in the group that received only antigen in the same period.

In order to determine the repeated dose toxicity and the amount of antibody production, vaccination of the rabbits was repeated 3 times (on days 0, 14, 28). On the 7th and 28th days after the last injection, the animals were bled, and serum samples were taken for antibody evaluation by indirect ELISA. Antibody titers were compared at a 1: 50,000 dilution of serum samples after the third and fifth injections. As illustrated in Figure 6, the data were very similar and there was no significant difference between these samples.

**Figure 6.**
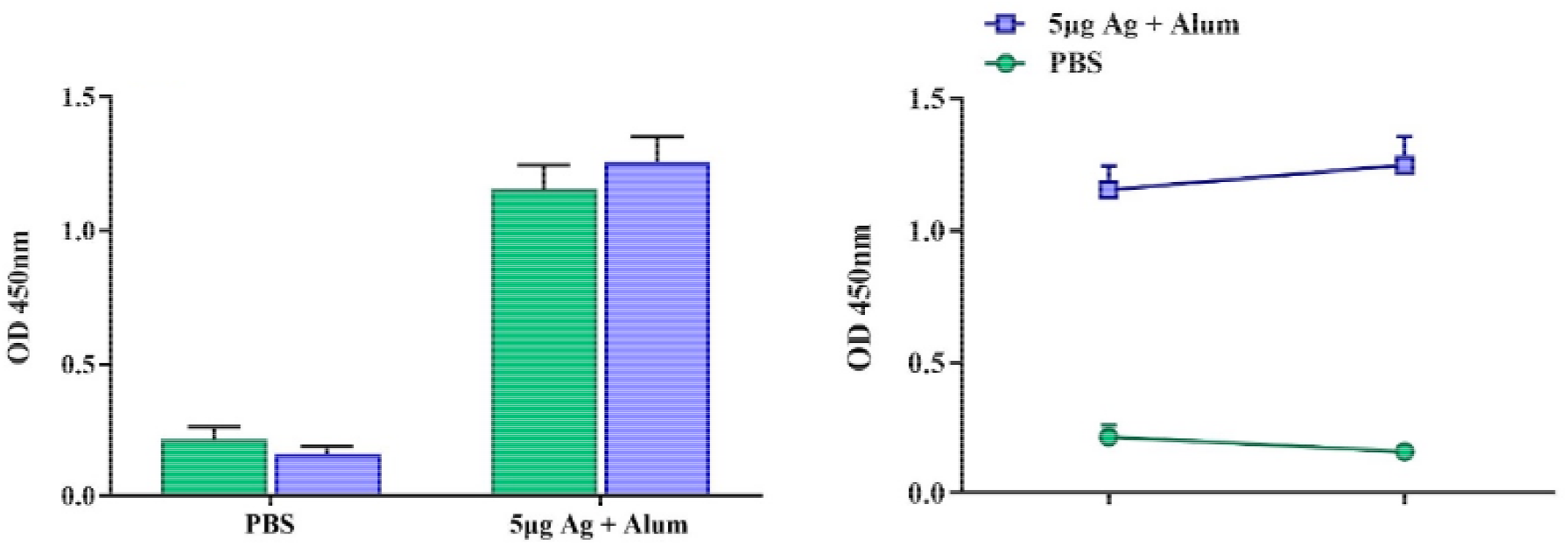
Induction of specific IgG antibodies by candidate vaccine in rabbit groups. Immunizations were carried out 3 times (on days 0, 14, 28) using 5 μg antigens in combination with alum. The control groups received PBS. After the third injections, the rabbits were bled and individual sera were assayed by ELISA at 7 (green) and 2 days (violet) after the last injection. Data are reported as mean ± SD.

Furthermore, the immunogenicity and protective efficacy of the COVIran Barekat vaccine were evaluated in Rhesus macaques, a nonhuman primate species that shows a COVID-19-like disease caused by SARS-CoV-2 infection.22 Macaques were immunized two times intramuscularly with 3 and 5 μg doses of COVIran Barekat vaccine on days 0 and 14. N and S-specific IgG antibody titer was determined over time in serum samples of monkey groups using ELISA. The control group did not show any detectable N and S-specific IgG antibody responses. However, N and S-specific IgG antibody titer emerged at week 1 and surged after 2 weeks in monkey groups that were immunized with the 3 μg or 5 μg doses of COVIran Barekat vaccine (antigen + alum). The titer of N and S-specific IgG was significantly (*P* < 0.05) higher than the control group (Figure 7).

**Figure 7.**
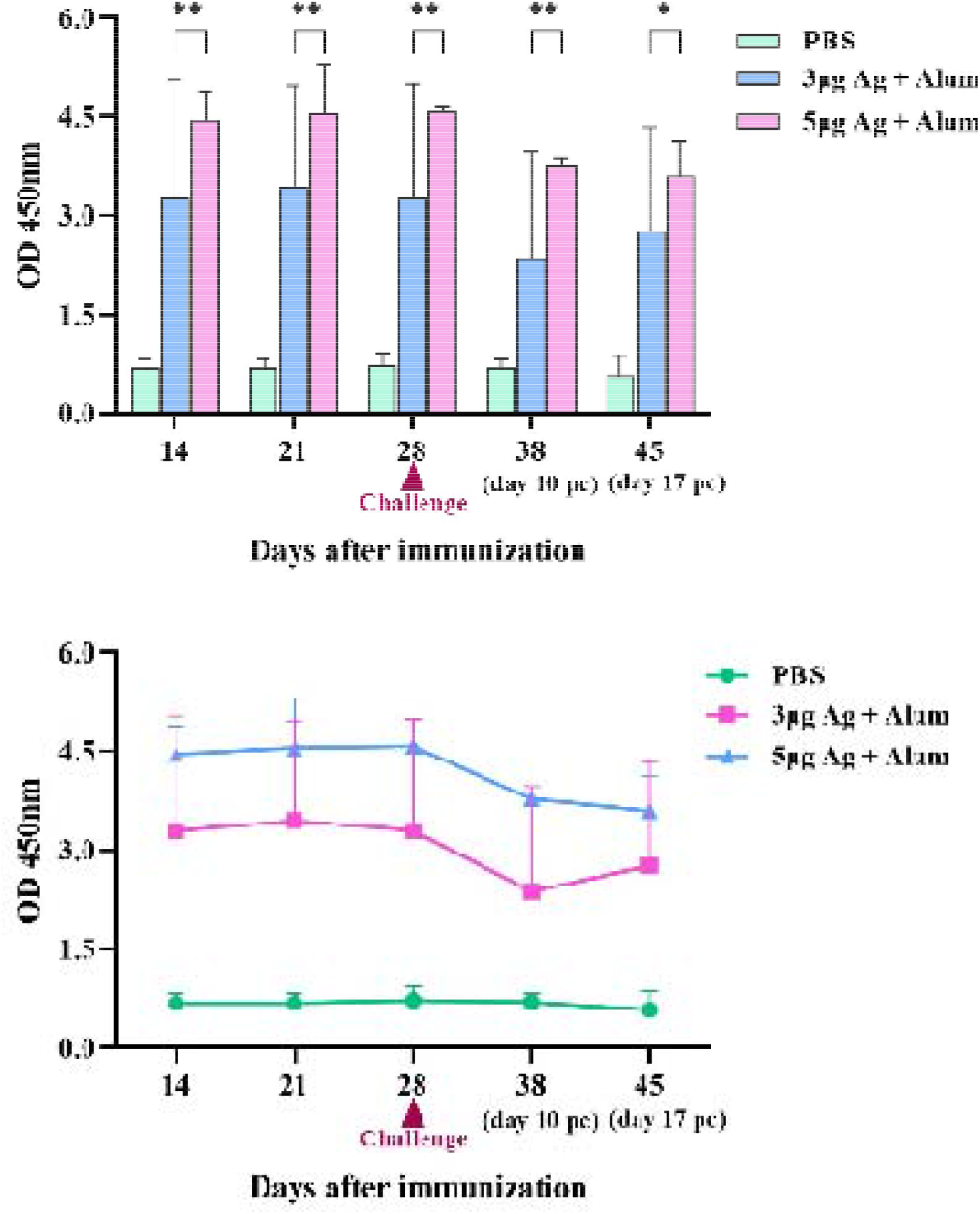
Induction of N and S-specific IgG antibodies by candidate vaccine in monkey groups. Rhesus macaques were intramuscularly immunized two times (at days 0 and 14) with 3 or 5 μg doses of candidate vaccine (antigen + alum). The control group was injected with PBS. The challenge study was carried out 10 days after the second immunization. N and S-specific IgG antibody titer in serum samples of monkey groups was determined by ELISA over time and data are reported as mean ± SD. The statistically significant differences between groups were determined using two-way ANOVA followed by Tukey’s post hoc test. Asterisks indicate significance: *P < 0.05 and **P < 0.001 in comparison with the control group.

The results of VNT in mice groups showed that the serum titer up to 1/512 has the ability to neutralize the virus while in rabbit groups, the serum titer up to 1/8192 has the ability to neutralize the virus which resulted after 5 injections. The virus neutralization antibody in monkey groups was 1/256.

Subsequently, a challenge study was conducted by direct inoculation of 10^4^ TCID50 of SARS-CoV-2 into the vaccinated and control macaques’ lungs intratracheally on day 28 (2 weeks after the second immunization) to verify the protective efficacy. All vaccinated macaques were highly protected against SARS-CoV-2 infection.

### b) Blood Biochemical and Hematological Analysis in Rhesus Macaques

A number of biochemical and hematological indicators were measured at different time points post-immunization. There was no significant difference in terms of biochemical markers in the vaccinated animals (Figure S1).

Hematological analysis after viral challenge in 3 µg vaccinated and unvaccinated animals showed lymphocytosis (Figure S2).

### c) Cytokines Assay in Lung Tissue of Rhesus Macaques

The lung tissue samples were used for protein extraction. The total protein concentration was determined by the Bradford assay. The cytokine assay was performed in homogenized tissue samples of cranial, middle, and caudal lung lobes by SDS-PAGE followed by western blot analysis using specific antibody and anti-β actin as a normalized control.

IL-6 (a pro-inflammatory cytokine), IL-4 (a Th2 lymphocytes marker), IL-10 (an anti-inflammatory mediator), and IFN-γ (an amplifier and stimulator cytokine in the immune system) were detected in the homogenized tissue samples of cranial, middle, and caudal lung lobes, using western blot and ELISA methods. The results of western blot showed that IL-6 decreased significantly in vaccinated animals compared to the control group (*P* < 0.0001). On the other hand, IL-4 and IL-10 increased significantly in 3 µg and 5µg vaccinated groups in comparison with the control group (P <0.05). Similarly, the results of ELISA showed that IL-4 and IFN-γ also increased significantly in vaccinated animals compared to the control group (P <0.05) (Figure 8).

**Figure 8.**
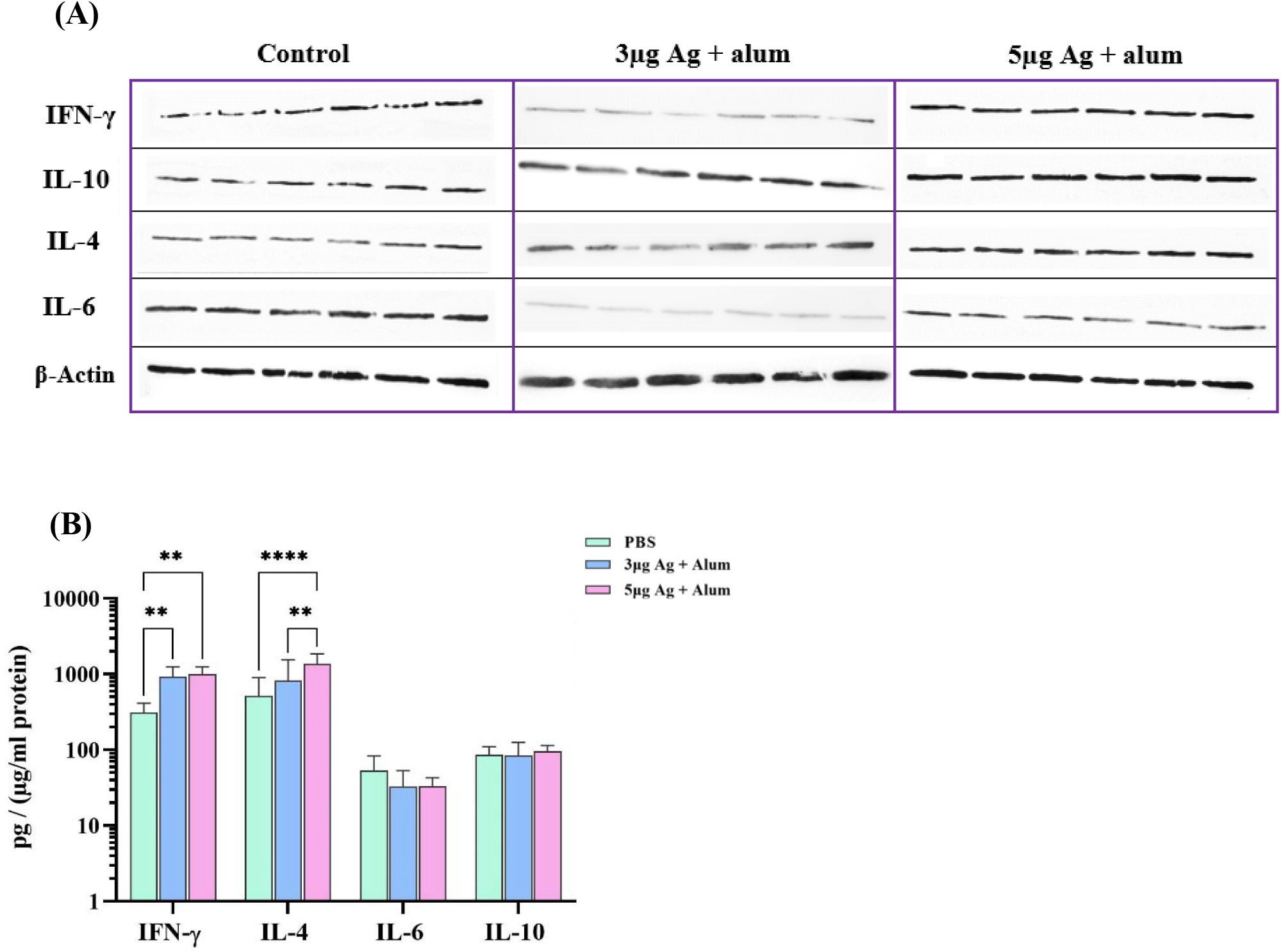
Cytokine level in homogenized lung tissue samples Macaques were immunized two times on days 0 and 14 through the intramuscular route with 3 μg or 5 μg antigen + alum. The control group was injected with PBS. A number of key cytokines in lung tissues were measured on day 45 after immunization by two methods (A and B) western blot analysis and (B) ELISA. The lung tissue samples were used for protein extraction. The total protein concentration was determined by the Bradford assay. Assessment of IL-6, IL-4, IL-10, IFN-γ in lung extracted proteins was performed by SDS-PAGE followed by western blot analysis, and also cytokine levels in lung tissue extracted total proteins were determined by ELISA. Cytokine levels were normalized to the total protein concentration in each lung homogenate. Data presented as mean ± SD. The statistically significant differences between groups were determined using two-way ANOVA followed by Tukey’s post hoc test. Asterisks indicate significance: *P < 0.05, ** P < 0.001, *** P < 0.0001 and **** P < 0.00001 in comparison with the control group.

## Evaluation of the Cellular Immune Response

### a) Granzyme B Activity Assay in Mice

The potential for induction of cellular immunity at different vaccine concentrations was evaluated by measurement of Granzyme B (Gzm B) activity using ELISA. The results revealed that Gzm B activity increased in the mice immunized with a mixture of antigen and alum adjuvant in comparison with the animals that received only antigen (*P* < 0.0001). Furthermore, there was a significant increase in Gzm B activity in the group that received the 5 μg dose of antigen + alum in comparison to the group which was immunized with the 3 μg dose of antigen + alum (Figure 9).

**Figure 9.**
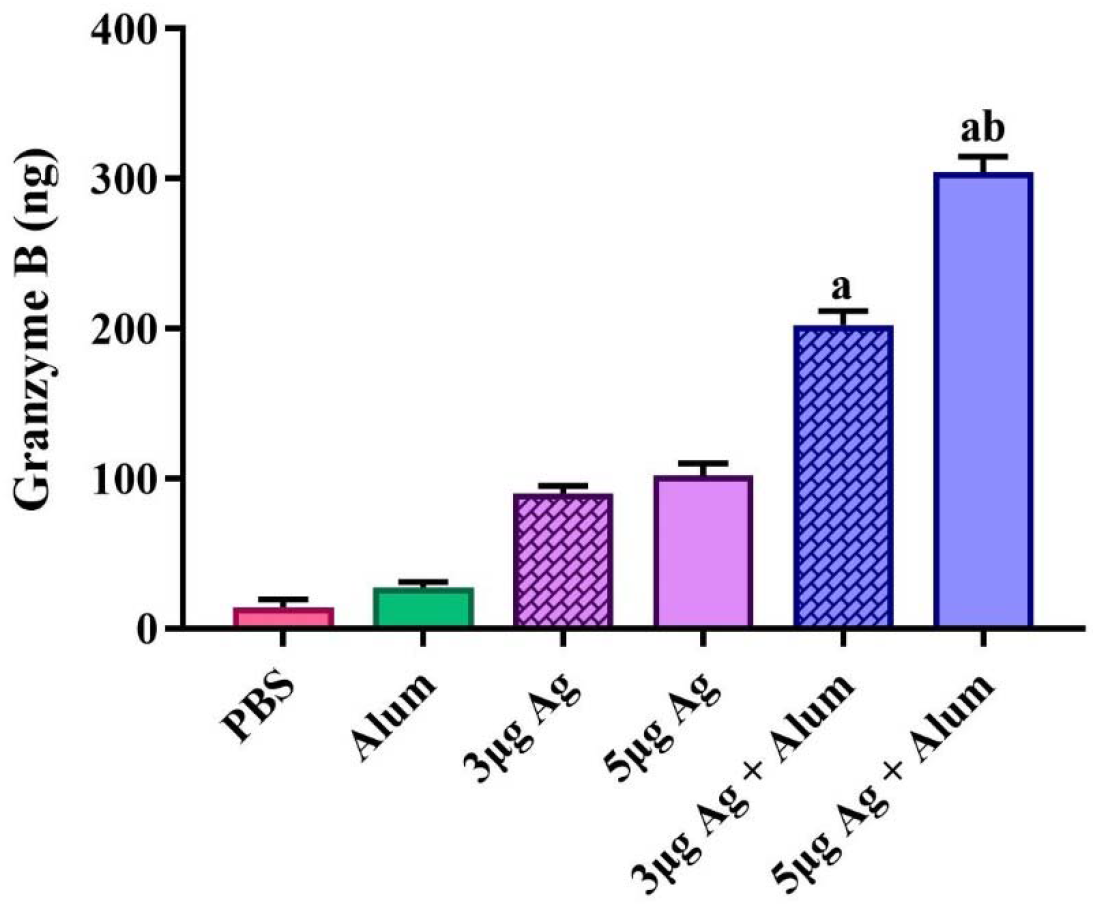
Measurement of Gzm B activity in mice groups that received different vaccine preparations. Immunizations were carried out two times over 2 weeks with 100 μL intramuscular injection of different vaccine preparations. The control groups received alum or PBS. At 2 weeks after the last immunization, spleens were isolated from immunized mice and Gzm B activity was measured. Data are reported as mean ± SD. The levels of statistical significance for differences between test groups were determined using one-way ANOVA followed by Tukey’s post hoc test. a: Statistical significance (P < 0.0001) in comparison with groups that received 3 and 5 μg antigen alone and b: Statistical significance (P < 0.0001) between groups that received 3 μg antigens with the groups immunized with 5 μg antigens.

### b) Viral Clearance from the Lungs of Immunized Rhesus Macaques after Challenge

Rhesus macaques injected with 3 μg candidate vaccine showed a significant reduction in lung virus titers compared with PBS-treated controls. The titer of virus was 10^2^ TCID50/mL in the group that received 3 μg while the PBS injected group had 10^6^ TCID50/mL. A higher level of viral clearance was detected in rhesus macaques immunized with 5 μg candidate vaccine and no viruses were detected.

### c) Histopathology and Immunohistochemistry

Histopathological evaluation of various organs, heart, spleen, liver, kidney, and brain in the vaccinated monkeys revealed no obvious lesion in the vaccinated animals. In infected control animals, in addition to severe lung lesions, renal tubular epithelium necrosis was found, probably due to SARS-CoV2. Histopathology of lung in infected control animals showed progressive moderate-to-severe interstitial pneumonia accompanied by macrophage and lymphocyte infiltration, particularly in the cranial and middle lobes. Moreover, we found moderate-to-severe thickening of alveolar septa with type II pneumocyte hyperplasia and fibrin deposition. Vasculitis with accumulation of lymphocytes around small vessels was detected in control monkeys’ lung tissue sections. In the vaccinated animals, the lung tissue showed diffuse mild interstitial pneumonia and mild bronchial epithelial necrosis limited to some areas in the cranial lobes (Figure 10).

**Figure 10.**
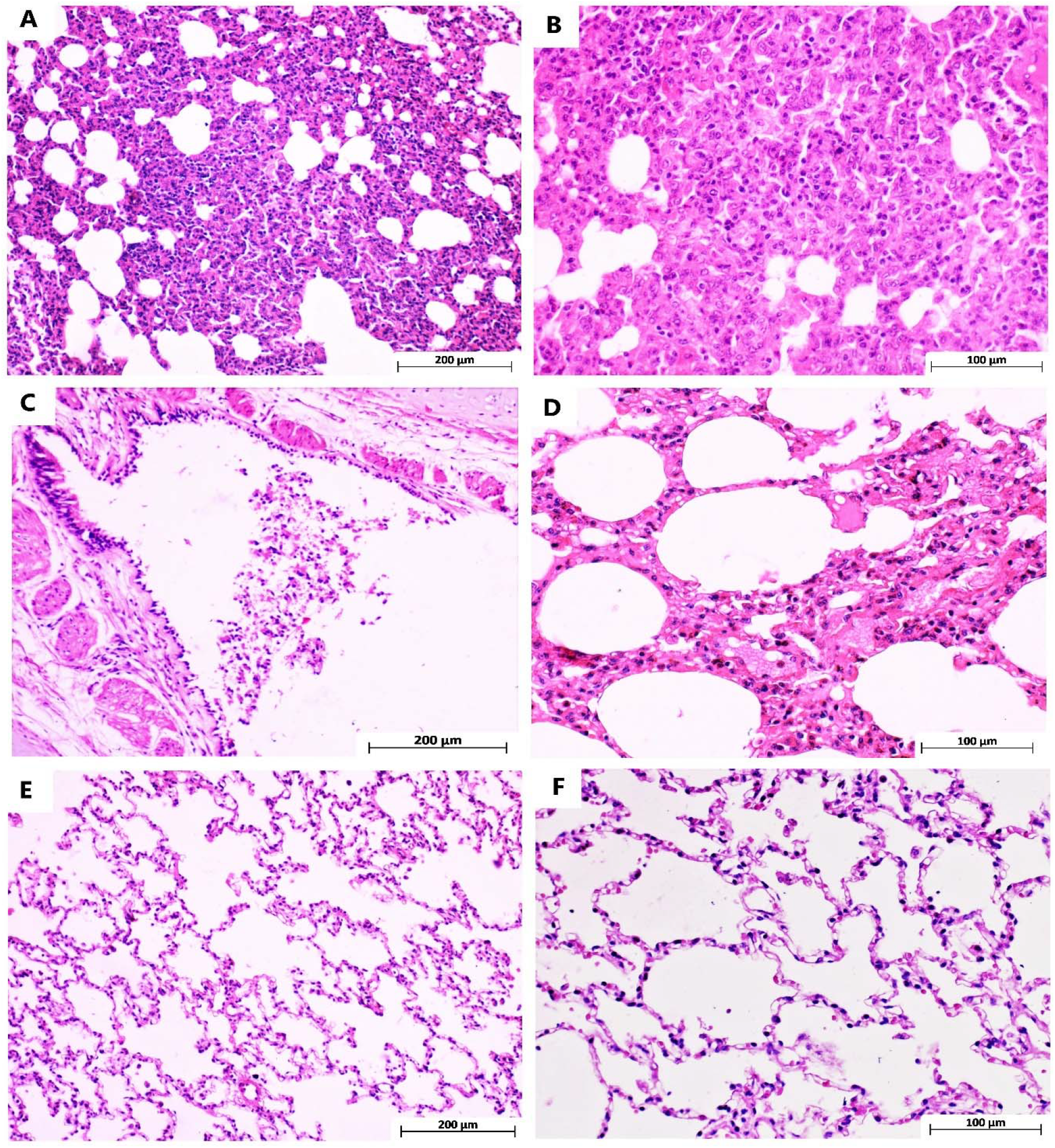
Mon keys ’ lung tissuee sections stained with hematoxylin and eosin. A and B are different magnifications of cranial and middle lung lobes in the control group, indicating severe interstitial pneumonia with marked thickening of alveolar septa and mononuclear inflammatory cell infiltration. C: bronchus section from the cranial lung lobe in the animal vaccinated with 5µg Ag+ alum which shows epithelial cell necrosis and sloughing due to SARS-Cov-2 infection. D: mild interstitial inflammation of only cranial lung lobe tissue in animals vaccinated with 3µg Ag+ alum. E and F show some areas of the cranial and the whole middle and caudal lung lobes in monkeys vaccinated with 3 and 5µg Ag+ alum, respectively.

Immunohistochemical characterization of immune cell population within the lung tissue sections revealed that the highest presence of CD68 positive M1 macrophages was detected in infected control monkeys. CD4 positive T lymphocytes were observed particularly near the vessels and airways of control monkeys’ lung tissue. More CD20 positive B cells and CD8 positive cytotoxic T cells were identified in the vaccinated animals compared to the control group (Figure 11).

**Figure 11.**
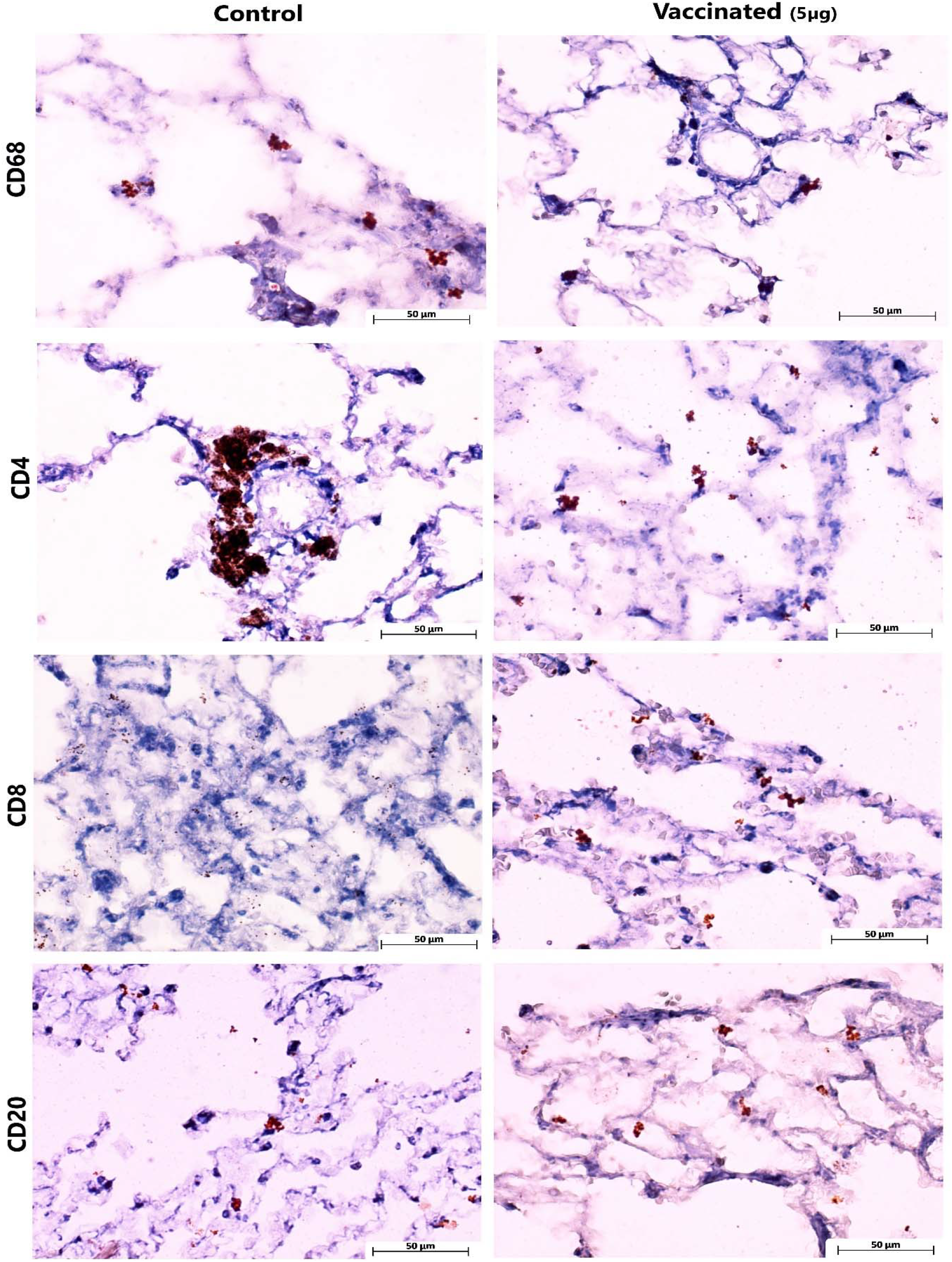
Immunohistochemical staining of immune cells within the lung tissue sections.

**Figure 12.**
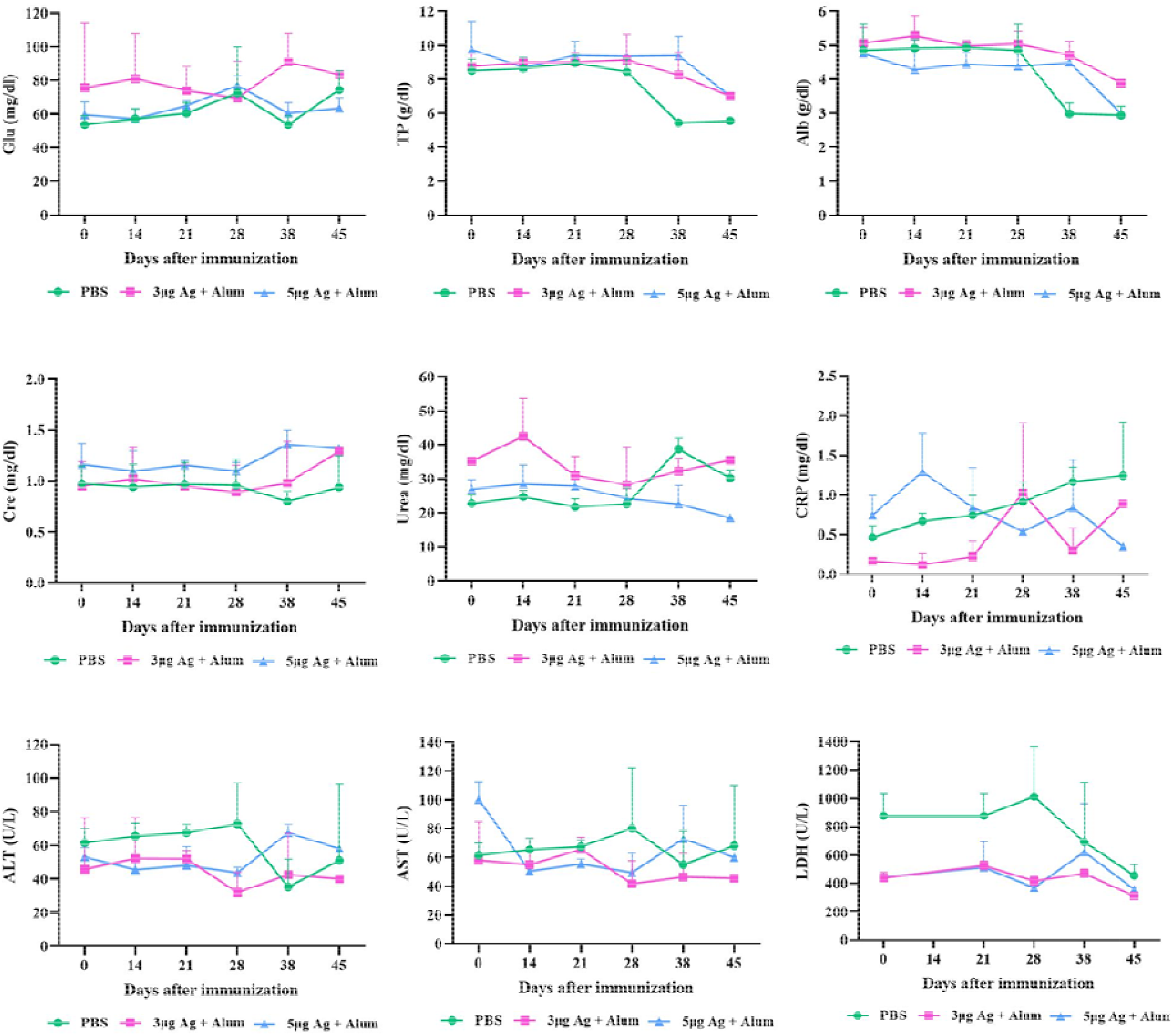
Biochemical indices are monitored to evaluate the safety of COVIran Barekat vaccine in Rhesus macaques. Macaques were immunized two times on days 0 and 14 through the intramuscular route with 3 μg or 5 μg antigen+ alum. Th control group was injected with PBS. A number of biochemical indices at different time points were measured by an animal calibrated autoanalyzer. Data points represent mean ± SD. Glu (Glucose), TP (Total protein), Alb (Albumin), Urea (Blood urea), Cre (Creatinine), CRP (C-reactive proteins), ALT (Alanine aminotransferase), AST (Aspartate aminotransferase), LDH (Lactat dehydrogenase). .

**Figure 13.**
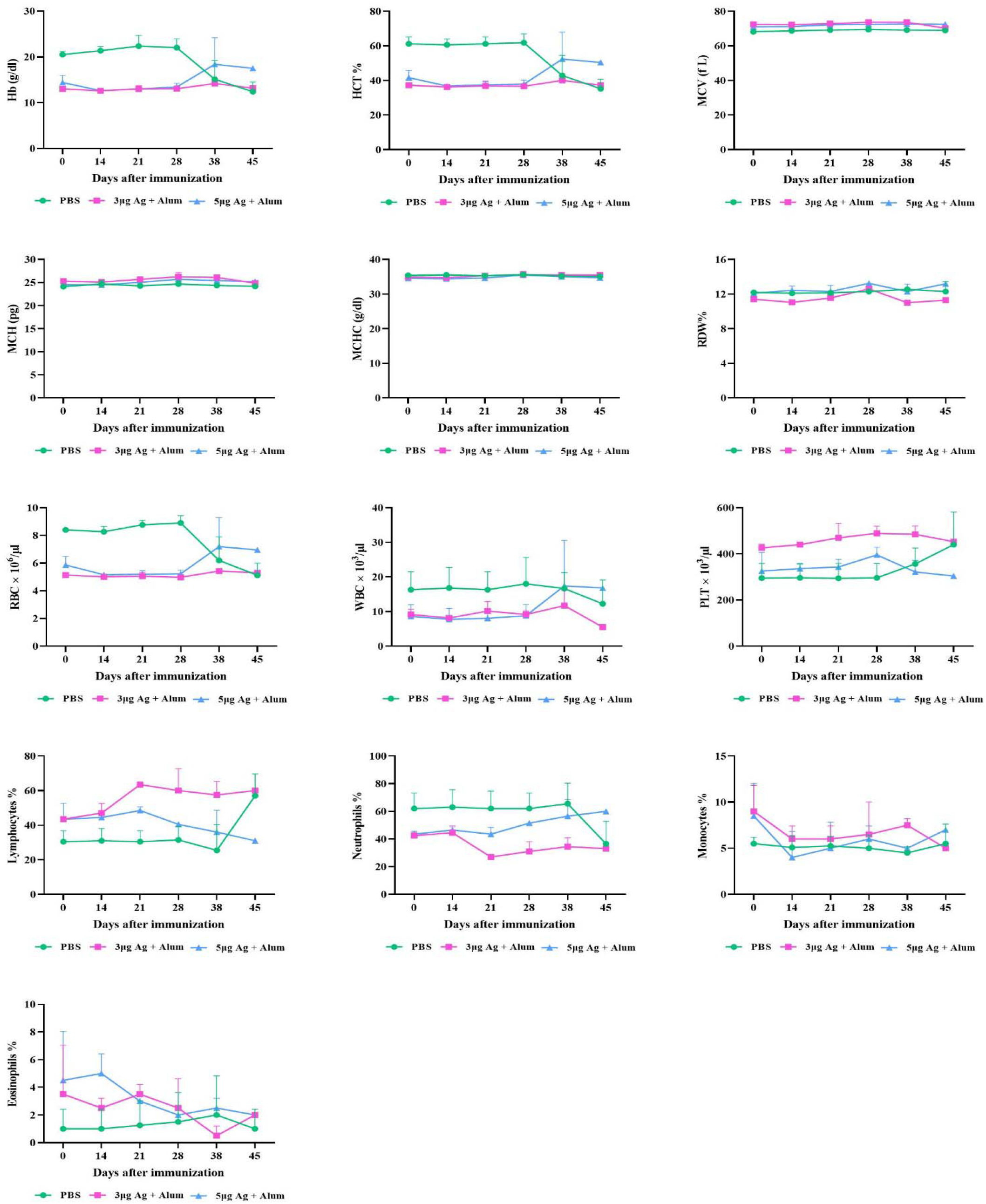
Hematological indices are monitored to evaluate the safety of COVIran Barekat vaccine in Rhesus macaques. Macaques were immunized two times on days 0 and 14 through the intramuscular route with 3 μg or 5 μg antigen+ alum. The control group was injected with PBS. A number of hematological indices were measured at different time points. Data points represent mean ± SD. Hb (hemoglobin), HCT (hematocrit), MCV (mean corpuscular volume), MCH (mean corpuscular hemoglobin), MCHC (mean corpuscular hemoglobin concentration), RDW (red cell distribution width), RBC (Red blood cell), WBC (white blood cell) and PLT (platelet count test)

## Discussion

Scientists have focused their efforts on the development of vaccines to stop the COVID-19 pandemic and its quick spread.23 To achieve this goal, several vaccine trials are being conducted worldwide. For decades, inactivated whole particle vaccines have been broadly administrated to prevent emerging respiratory diseases.24 The results from preclinical and clinical studies of three inactivated COVID-19 vaccines have shown promising outcomes against SARS-CoV2.25-28

On the other hand, the application of adjuvants in vaccine formulation can increase and/or modulate the intrinsic immunogenicity of antigen and elicit strong and long-lasting immune responses.29 It may also substantially reduce the amount of antigen and/or the number of boost injections required to achieve the desired immune responses.30

In the current study, SARS-CoV-2 was isolated from the nasopharyngeal sample of a COVID-19 infected patient. Viruses were isolated and cultured in Vero cells, then inactivated with β-propiolactone for 24 hours. Then, they were concentrated via ultrafiltration and purified with sucrose gradient ultracentrifuge or Ion-exchange Chromatography (IEC) and Size Exclusion Chromatography (SEC). The purified viruses were sterilized through filtration and were formulated with Alum adjuvant to serve as the COVIran Barekat SARS-CoV-2 candidate vaccine. After safety assays, it was injected into rodent and non-rodent animal models to evaluate the titer of antibody level. In addition, Rhesus macaques were immunized with two doses of the vaccines and challenged intratracheally with 10^4^ TCID50 of SARS-CoV-2.

Measurement of anti-COV-2 IgG antibody titers in the serum of vaccinated mice showed that the injection of the candidate vaccine (purified inactivated viruses in combination with adjuvant) significantly induced a high level of specific antibodies on day 21 and day 42 after the first injection.

The administration of alum adjuvant causes the high antibody response to persist for 42 days after immunization due to the continuous release of antigen. These results also show that the antibody produced has adequate durability after 42 days of vaccination and can neutralize the wild virus. Therefore, alum adjuvant is a good option for vaccine formulation and enhancement of the immune response.

The VNT was performed using different dilutions of sera in vaccinated mice and rabbits after the second and third injections, respectively. The results showed that the elevated antibodies have high affinity to neutralize the virus in cell culture.

The immunization of Rhesus macaques with the two-dose COVIran Barekat vaccine led to efficient protection against 10^4^ TCID50 of SARS-CoV-2 intratracheal challenge compared to controls.

One concern about COVID-19 vaccines is the antibody-dependent enhancement (ADE) phenomenon, whereby the vaccine could make the subsequent SARS-CoV-2 infection more severe.24,31 The ADE phenomenon has been reported in studies of Middle East respiratory syndrome–CoV and SARS-CoV vaccines in animal challenge models.32 However, this effect was not observed in an immunization-challenge model of Rhesus macaques using the same vaccines in the preclinical study or in the reports from preclinical studies of other inactivated COVID-19 vaccine candidates.25,33 Studies have shown that previous infection with SARS-CoV-2 could protect against re-challenge in Rhesus macaques. The ADE phenomenon may happen due to an imbalance between the T helper 1 (Th1) and T helper 2 (Th2) responses.

Granzyme B (Gzm B), a neutral serine protease, is expressed exclusively in the granules of activated cytotoxic T lymphocyte (CTL), natural killer cells, and lymphokine-activated killer cells. Gzm B induces CTL-mediated DNA fragmentation and apoptosis in allogeneic target cells.34 Gzm B activity assays can be used for studying T cell immunity, CTL responses, cell-mediated cytotoxicity, and apoptosis.35 The efficacy of vaccines in the production of virus-neutralizing antibodies (VNA) and activation of Gzm B was investigated. The results showed that the candidate vaccine could induce high VNA level and Gzm B activity.

Another concern associated with whole-inactivated virus-based vaccines, particularly those with the alum adjuvant that can induce T helper 2 cell–based responses, is the probability of vaccine-associated enhanced respiratory disease (VAERD). VAERD was reported in young children in the 1960s when the whole-inactivated virus vaccine with alum adjuvant was tested for measles and respiratory syncytial virus.36,37 However, most of the inactivated vaccines under development against COVID-19 have used alum adjuvant, and no evidence of VAERD has been observed yet. On the contrary, alum may reduce immunopathology compared with un-adjuvanted coronavirus vaccines.14,38,39 Also, another assertion that aluminum-adjuvanted vaccines induce autism or other chronic illnesses has been thoroughly discredited.

**In conclusion**, taken together, data obtained from a preclinical study of COVIran Barekat candidate vaccine shows adequate safety and efficiency and thus, it may be a promising and feasible prophylactic vaccine to confer protection against the SARS-CoV-2 infection. Also, alum would serve as an effective adjuvant in the formulation of SARS-CoV2 and other candidate vaccines. These results suggest a path forward to Phase I, II, and III clinical trials with the COVIran Barekat vaccine.

